# Mendelian randomization analysis revealed causal effects from gut microbiota to abdominal obesity

**DOI:** 10.1101/2020.04.20.052407

**Authors:** Qian Xu, Shan-Shan Zhang, Yu-Fang Pei, Jing-Jing Ni, Lei Zhang, Rui-Rui Wang, Yu-Jing Weng, Xun Cui, Xin-Tong Wei

## Abstract

Although recent studies have revealed the association between the gut microbiota and obesity, the causality remains elusive. We performed a Mendelian Randomization (MR) analysis to determine whether there is a causal relationship between gut microbiota and abdominal obesity. We used a two-sample MR approach to assess the causal effect from gut microbiota to obesity based on genome-wide association studies (GWAS) summary statistics. The GWAS summary statistics of gut microbiota obtained from UK-twins cohort (N=1,126) were used as discovery sample exposure, and the GWAS summary statistics from the Genetic Environmental Microbial (GEM) project (N=1,098) were used as replication sample exposure. Trunk fat mass (TFM) summary statistics from the UK Biobank (UKB) cohort (N=330,762) were used as outcome. Bacteria were grouped into taxa features at family level. A total of 16 families were analyzed in the discovery sample. Family *Barnesiellaceae* was associated with TFM at the nominal significance level (*b*=-3.81×10^−4^, *P*=1.96×10^−3^). The causal association was successfully replicated in the replication sample (*b*=-7.34×10^−3^, *P* =2.77×10^−2^). Our findings provided evidence of causal relationship from microbiota to fat development, and may be helpful in selecting potential causal bacteria for manipulating candidate gut microbiota to therapy obesity.

**IMPORTANCE:** Obesity, as a global public health problem, is one of the most important risk factors contributing to the overall global burden of disease, and is associated with an increased risk of cardiovascular disease, type 2 diabetes, and certain cancers. Recent studies have shown that gut microbiota is closely related to the development of obesity, but the causal relationship is unclear. Therefore, it is necessary to identify the causality between gut microbiota and obesity. The significance of our research is in identifying the causal relationship from specific bacteria to fat development, which will provide the new insights into the microbiota mediated the fat development mechanism.

## INTROUDCTION

Obesity is a chronic metabolic disease characterized by excessive accumulation of adipose tissue. It is one of the most important risk factors contributing to the overall burden of diseases worldwide, associated with increased risk of cardiovascular disease, type 2 diabetes and certain cancers(1). In 2013, the number of overweight and obese individuals globally has reached 2.1 billion and the prevalence has been increasing substantially(2).

Body mass index (BMI), which is defined as body mass in kilograms divided by the square of height in meters (kg/m^2^), is currently the standard measure of obesity due to its simplicity. However, BMI is never the ideal phenotype to measure obesity because it does not give a precise idea about the body composition(3). Human body mass is composed of fat mass, lean mass, bone mass, water and soft tissues; it is only fat mass that induces obesity and causes a series of adverse clinical manifestations. Therefore, fat mass is the only accurate and ideal phenotype to measure obesity(4, 5). Nonetheless, the research using fat mass as a measure of obesity has rarely been studied. Among various types of fat-induced obesity, abdominal obesity is perhaps the most severe. Fat stored in the abdomen is more harmful than fat stored at other body regions. For example, fat mass stored more centrally leads people to be more susceptible to cardiovascular diseases and endocrine disorders(6).

Even though obesity can be attributable to lifestyle, culture factors and genetics(7–9), mounting evidence demonstrated that the human gut microbiome play an important role in the development of obesity(10–12). Mice models provide the causal evidence of obesity linked to gut microbiome, but the finding are far from consistent(13, 14). A case-control study found the abundance of *Lactobacillus reuteri* was positively correlated with BMI, and *Bifidobacterium animalis, Methanobrevibacter smithii*, and *Escherichia coli* were negatively associated with BMI(15). A cohort study identified 34 bacterial taxa associated with BMI and explained 4.5% of its variance(16). Nonetheless, the causality between specific taxa of gut microbiota and obesity is still ambiguous due to many confounding factors (including lifestyle, diet and disease status) that occur within the population.

Mendelian randomization (MR) analysis is a statistics approach that uses genetic variants as instrumental variables (IVs) to test the causality from potentially risk exposure to health outcomes in a cross-sectional study. It is less likely to be affected by confounding factors or inverse causation than conventional observational studies(17, 18). Previous study has shown that host genetic variations influence the composition of gut microbiota(19). Recent years, increasing genome-wide association studies (GWAS) for gut microbiota(20–24) make it possible to infer causal relationship by performing MR analysis base on summary statistics of GWAS.

In the present study, we conduct a two-sample MR study(25) to investigate the causal link from specific taxa of gut microbiota to trunk fat mass (TFM) using summary statistics of GWAS. Specifically, the summary statistics from microbial GWAS serve as exposure while the summary statistics from trunk fat mass GWAS serve as outcome.

## RESULTES

In the discovery TwinsUK sample, there are total of 229 SNPs associated with gut microbiota at the significance level *P*<1.0×10^−5^. After clumping, there were 102 SNPs, categorized into 16 bacteria families (**Supplementary table 1**). The family with the largest number SNPs is *Ruminococcaceae* (24 SNPs), followed by *Lachnospiraceae* (23 SNPs) and *Bacteroidaceae* (21 SNPs). There were 6 families each containing only one SNP, *Bifidobacteriaceae, Streptococcaceae, Veillonellaceae*, *Barnesiellaceae*, *Enterobacteriaceae* and *Porphyromonadaceae*. The number of IV SNPs ranged from 2 to 6 for the remaining 7 families.

To ensure that the above IVs are free from horizontal pleiotropy, we performed MR-PRESSO analysis on independent SNPs to detect the potential SNPs with pleiotropy effect. One out of 6 IVs in family *Clostridiaceae*, 1 out of 21 IVs in family *Bacteroidaceae*, 3 out of 23 IVs in family *Lachnospiraceae*, 4 out of 24 IVs in family *Ruminococcaceae* and 1 out of 6 IVs in family *Pasteurellaceae* were detected as outliers using the MR-PRESSO outlier test (**Supplementary Table 2**).

After removing the SNPs with pleiotropy effect, we performed MR analysis on the remaining SNPs. In the discovery sample, only one family *Barnesiellaceae* is nominally significant level (*beta*=-3.81×10^−4^, *P*=1.96×10^−3^). Specifically, this family *Barnesiellaceae* contains only one IV SNP rs4897946, which is located in the intron region of *MIER2* gene on chromosome 19 (**Table 1**).

**Table 1.**
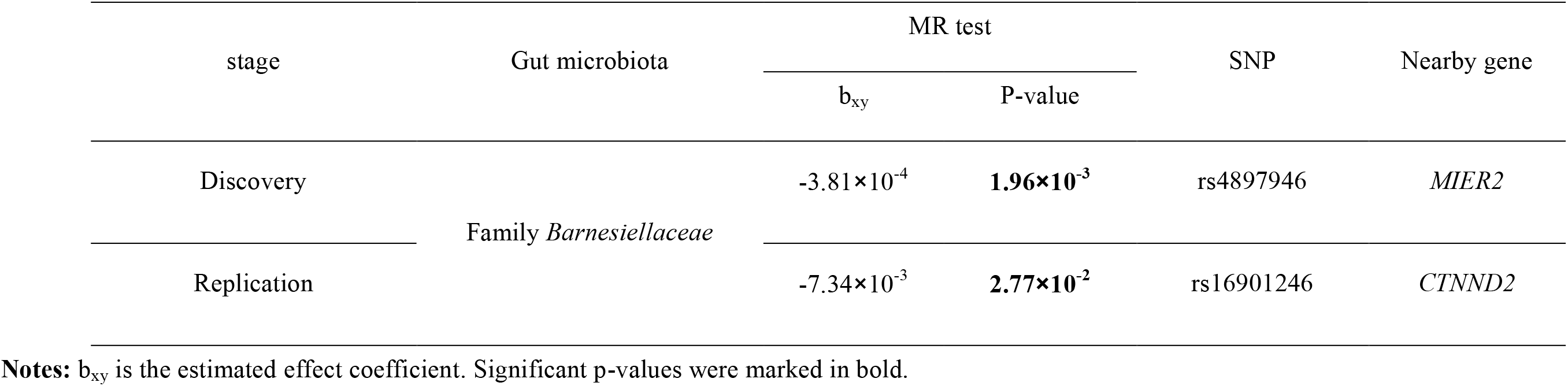
Causal estimations of gut microbiome on trunk fat mass in the discovery and replication cohorts

| stage | Gut microbiota | MR test |  | SNP | Nearby gene |
| --- | --- | --- | --- | --- | --- |
| | | $b_{xy}$ | P-value | | |
| Discovery | Family *Barnesiellaceae* | $-3.81 \times 10^{-4}$ | **$1.96 \times 10^{-3}$** | rs4897946 | *MIER2* |
| Replication | | $-7.34 \times 10^{-3}$ | **$2.77 \times 10^{-2}$** | rs16901246 | *CTNND2* |

**Notes:** $b_{xy}$ is the estimated effect coefficient. Significant p-values were marked in bold.

The significant family *Barnesiellaceae* is subjected to be replicated in the GEM replication sample. Again, only one SNP rs16901246 is assigned to this family. Interestingly, both the causal effect direction (*beta*=-7.34×10^−3^) is consistent with that in the discovery sample and the p-value is significant (0.03), strengthening the confidence towards the true association of this family. The IV SNP rs16901246 is located in the intron region of *CTNND2* gene on chromosome 5.

## DISCUSSION

In this study, we performed a two sample MR-based causality analysis between gut microbiota and TFM using summary statistics from GWAS summary statistics. By combining the results from discovery and replication studies, we identified a causal association from bacteria family *Barnesiellaceae* to TFM. Specifically, our results demonstrated a reverse causal effect from the former to the latter.

The gut microbiota of healthy adult was primarily dominated by two phyla *Firmicutes* (53.9% of total) and *Bacteroidetes* (35.3%), with other phyla including *proteobacteria, Verrucomicrobia, Actinobacteria*, and *Tenericutes*(26, 27). Previous studies have shown the relative abundance of *Firmicutes* and *Bacteroidetes* in obese populations. For example, a twins study revealed that the proportion of *Bacteroidetes* is higher in obese compared with lean individuals(12). Another animal study found a reduction in the abundance of *Bacteroidetes* together with a relative increase in *Firmicutes* in obese animals compared with lean animals(13). The family *Barnesiellaceae* identified in the present study is a member of *Bacteroidetes* phylum. A recent study found that the family *Barnesiellaceae* was correlated with the percentage of body fat and modified by exercise(28). In a case-control study, Chierico et al reported the abuandance of family *Barnesiellaceae* may be a microbial biomarker in healthy adolescents(29). These previous observational studies provide valuable clues towards the close relationship between *Barnesiellaceae* and fat mass development. For the first time, to our best knowledge, the present study established a causal link from the former to the latter.

A possible mechanism of gut microbiota influence the development of obesity is that gut microbiota can increase energy production from diet, contribute to low-grade inflammation and regulate fatty acid tissue composition(30). Though it remains unclear for the mechanism underlying the regulator path from *Barnesiellaceae* to obesity developement, previous study showed that the *Barnesiellaceae* has been associated with low-fiber consuming(31). Another study found the relative abundance of *Barnesiellaceae* clearly decreased in a medium containing only proteins and peptones, which revealed it not involve in protein breakdown and fermentation(32). However, further functional investigation is warranted to validate this correlation.

The MR approach is robust to confounding factors and reverse causality in observational studies (33). In this study, we applied a two sample MR approach based on summary statistics to explore the causal relationship between gut microbiota and TFM. Our study has following advantages. First, it is based on large-scale GWAS summary statistics that are publicly available, thus offers an efficient option to mine reliable genetic information without additional experiment costs. Second, we used TFM instead of BMI as a phenotype to measure abdominal obesity, which provided exactly accurate risk information of obesity.

However, there are also some limitations in our study. Firstly, the gut microbiota GWAS is still scarce, resulting in very limited t gut microbiota-associated SNPs to be used for analysis. Secondly, the significant causal association identified in this study were obtained using single IV, which has inferior robustness and statistical power.

In conclusion, by performing a two sample MR analysis based on several GWAS summary statistics, we identified a causal relationship from gut microbiota to abdominal obesity. Our results may be helpful in selecting potential causal bacteria for manipulating candidate gut microbiota to therapy obesity.

## MATERIALS AND METHODS

### Ethics statement

Gut microbiota GWAS summary statistics were accessed from published studies. No new IRB approval was required.

Trunk fat mass sample came from the UKB cohort, which is a large prospective cohort study of ~500,000 participants from across the United Kingdom, aged between 40-69 at recruitment. Ethics approval for the UKB study was obtained from the North West Centre for Research Ethics Committee (11/NW/0382), and informed consent was obtained from all participants. This study (project number 41542) was covered by the general ethical approval for the UKB study.

### GWAS summary statistics for gut microbiota

For exposure, we collected publicly available GWAS summary statistics of gut microbiota from two independent studies: the TwinsUK study and the Canada Genetic Environmental Microbial (GEM) project study. The TwinsUK study was used as discovery sample and it consisted of 489 dizygotic (DZ) twin pairs and 637 monozygotic (MZ) twin pairs with an age range of 18-89 years(22). The GEM project was used as replication sample, which included 1,098 healthy first-degree relatives of patients with Crohns disease between 6 and 35 years of age (24). Stool collection, DNA extraction, 16 sRNA gene sequencing and taxa filtering were performed on both cohorts.

In the discovery sample, the genetic associations between 945 bacteria taxa and 1,300,091 host SNPs were tested. A total of 307 host SNPs were identified to be associated with 61 bacteria taxa (1 kingdom + 6 phyla + 9 classes + 9 orders + 16 families + 16 genera + 4 species) at a FDR<0.2. The *P* values at these SNPs ranged from 4.94×10^−9^ to 7.33×10^−5^. The summary statistics of these significant SNPs were assessed through the supplemental table of the study publication(22).

In the replication sample, the associations between 3,727,707 host SNPs and 166 non-redundant bacterial taxa were examined. A total of 58 SNPs were identified to be associated with the relative abundance of 33 taxa at the genome-wide significance level (*P*<5×10^−8^). The summary statistics of these significant SNPs were assessed through the supplemental table of the study publication(24).

### UKB trunk fat mass sample

All the included participants in the UKB sample are those who self-reported as white (data field 21000). Participants who had a self-reported gender inconsistent with the genetic gender, who were genotyped but not imputed or who withdrew their consents were removed.

Trunk fat mass (TFM) was measured by bioelectrical impedance analysis approach. Phenotypic outliers were monitored by the Tukey method. Covariates, including age, sex, assessment center (23 levels), genotyping batch (2 levels) and the top 10 principal components (PCs) derived from genome-wide genotype data, were used to adjust raw phenotype. The residuals were normalized into inverse quantiles of standard normal distribution, which were used for subsequent association analysis.

Genome-wide genotypes were available for all participants at 784,256 genotyped autosome markers, and were imputed into UK10K haplotype, 1000 Genomes project phase 3 and Haplotype Reference Consortium (HRC) reference panels. A total of ~92 million variants were generated by imputation.

We used BOLT-LMM to perform linear mixed model (LMM) analysis of genetic association. The LMM analysis can adjust for population structure and relatedness.

### Genetic instrumental variants selection

Based on the GWAS summary results of gut microbiota,a series of quality control (QC) criteria were applied to select eligible genetic instrumental variables (IVs). Specifically, bacteria taxa were analyzed at the family level. A feature was defined as a distinct family. For SNP with multiple signals within one feature, the strongest signal was selected for that feature.

In the discovery sample, SNPs associated with microbiota at the α=1×10^−5^ level were selected and assigned into distinct features. The SNPs within each feature were then clumped with PLINK (v1.9) to retain independent SNPs only. For clumping The linkage disequilibrium (LD) threshold was set to be *r*^2^<0.1 and the clumping window size was set to be 500 kb, where LD was estimated based on the 1000 genomes project sequencing data (phase 3).

In the replication sample, SNPs of association at the same α=1×10^−5^ were not accessible. In contrast, only SNPs significant at the α=5×10^−8^ level were reported. Therefore, all the reported SNPs were selected. SNPs were again assigned into features and clumped to retain independent SNPs, following the same steps as those used in the discovery sample.

### Removal of horizontal pleiotropy

We applied the MR-PRESSO Global test(34) and Outlier test to detect potential horizontal pleiotropy. The MR-PRESSO global test evaluates overall horizontal pleiotropy among all SNPs, and the MR-PRESSO Outlier test evaluates the presence of specific horizontal pleiotropic outlier variants by calculating the p-value of each SNP pleiotropy significance. The MR-PRESSO global test was first applied to evaluate overall pleiotropy. In the presence of pleiotropy, the MR-PRESSO Outlier test was then applied and the SNP with the smallest pleiotropy p-value was removed. The MR-PRESSO Global test was again performed on the remaining SNPs. The process repeated until the Global test was non-significant (*P*>0.05).

The final retained SNPs were used as non-pleiotropic IVs to perform subsequent Mendelian randomization analysis.

### Mendelian randomization analysis

We performed two sample MR analysis to examine the causal effect from bacteria taxa to TFM. Specifically, we tested the association of the identified IVs within each bacteria taxa with TFM. For bacteria taxa containing multiple SNPs, we used five methods to estimate the causal effect, including the inverse variance weighted (IVW) test(35), the MR-Egger regression(36), the weighted median estimator(37), the simple mode-based estimator and the weighted mode-based estimator(38). The results were mainly based on the IVW method while the other 4 methods served as its complement. For bacteria taxa containing only one SNP, the Wald Ratio method was used for MR analysis. This method calculates the causal effect by using the coefficient of the SNP-outcome association divided by the coefficient of the SNP-exposure association(39).

Significant families identified in the discovery TwinsUK study were subjected to be replicated in the replication GEM study, following the same MR analysis procedure.

All the above analyses were performed with the R packages *TwoSampleMR* (https://github.com/MRCIEU/TwoSampleMR)(40) and *MR-PRESSO* (https://github.com/rondolab/MR-PRESSO)(34)

## ACKNOWLEDGMENT

This research was conducted using the UK Biobank resource under application number 41542. We appreciate all the volunteers who participated in this study. We are grateful to the TwinsUK study and the Genetic Environmental Microbial (GEM) project for releasing the gut microbiota GWAS summary statistics.

YFP and LZ are partially supported by the funding from national natural science foundation of China (31771417 and 31571291) a project funded by the Priority Academic Program Development (PAPD) of Jiangsu higher education institutions and the Undergraduate Training Program for Innovation and Entrepreneurship, Soochow University (201810285048Z). The numerical calculations in this paper have been done on the supercomputing system of the National Supercomputing Center in Changsha.

